# The geometry of photopolymerized topography influences neurite pathfinding by directing growth cone morphology and migration

**DOI:** 10.1101/2023.08.28.555111

**Authors:** Joseph T. Vecchi, Madeline Rhomberg, C. Allan Guymon, Marlan R. Hansen

## Abstract

Cochlear implants (CIs) provide auditory perception to those with profound sensorineural hearing loss: however, the quality of sound perceived by a CI user does not approximate natural hearing. This limitation is due in part to the large physical gap between the stimulating electrodes and their target neurons. Therefore, directing the controlled outgrowth of processes from spiral ganglion neurons (SGNs) into close proximity to the electrode array could provide significantly increased hearing function. For this objective to be properly designed and implemented, the ability and limits of SGN neurites to be guided must first be determined. In this work, we engineered precise topographical microfeatures with angle turn challenges of various geometries to study SGN pathfinding. Additionally, we analyze sensory neurite growth in response to topographically patterned substrates and use live imaging to better understand how neurite growth is guided by these cues. In assessing the ability of neurites to sense and turn in response to topographical cues, we find that the geometry of the angled microfeatures determines the ability of neurites to navigate the angled microfeature turns. SGN neurite pathfinding fidelity can be increased by 20-70% through minor increases in microfeature amplitude (depth) and by 25% if the angle of the patterned turn is made more obtuse. Further, by using engineered topographies and live imaging of dorsal root ganglion neurons (DRGNs), we see that DRGN growth cones change their morphology and migration to become more elongated within microfeatures. However, our observations also indicate complexities in studying neurite turning. First, as the growth cone pathfinds in response to the various cues, the associated neurite often reorients across the angle topographical microfeatures. This reorientation is likely related to the tension the neurite shaft experiences when the growth cone elongates in the microfeature around a turn. Additionally, neurite branching is observed in response to topographical guidance cues, most frequently when turning decisions are most uncertain. Overall, the multi-angle channel micropatterned substrate is a versatile and efficient system to assess SGN neurite turning and pathfinding in response to topographical cues. These findings represent fundamental principles of neurite pathfinding that will be essential to consider for the design of 3D systems aiming to guide neurite growth *in vivo*.

## 1. Introduction

Cochlear implants (CIs) are prosthetics that replace the mechanosensory transduction of sound by directly stimulating spiral ganglion neurons (SGNs) to provide hearing sensation. Tremendous improvements have been made to neural electrode devices in recent years, but they fail to fully emulate the sensory functions they seek to restore.[1,2,3] In particular, CIs are limited by the number of independent perceivable channels they provide.[4,5] Much of the unrealized potential of this device arises from poor tissue integration, with a large distance between the stimulating electrodes and their target, SGNs in the spiral ganglion.[6,7,8] CI electrode arrays are typically inserted in the scala tympani and reside hundreds of microns away from the SGN somatas in the ganglion, compared to the tens of nanometer gap of the synapse it is seeking to functionally replace. Enhancing the neural – electrode interface by reducing this gap is an essential element of next generation neural prostheses, as they seek to recapitulate the architecture and function of native neural systems more accurately.[9,10,11,12]

To improve the ability of neural prostheses to **mimic native neural function**, there is intense interest in developing technologies that could guide axon regeneration into close proximity to the stimulating electrode.[11,13] Further, this aspiration is not limited to SGNs as others are working to direct the growth of other neurons such as dorsal root ganglion neurons (DRGNs). To inform these aims, many are studying how neurites grow and turn in response to various cues such as diffusible chemical gradients,[10,14] patterned peptide surface coatings,[15,16] and engineered surface topography,[17,18] among others.[19] Though each of these approaches has its optimal uses and limitations, engineered surface topography to direct cell material interactions are particularly appealing for directing neurite growth to improve neural prostheses.

First, topographical cues offer tremendous control of cell behavior with precise modifications and reproducibility. These engineered features can be controlled with submicron resolution and the pattern geometry (periodicity, amplitude/depth, and shape) can be easily modulated to explore the relationship between the topography and neurite guidance.[17] Additionally, methacrylate polymer systems are shelf stable and are used extensively in a variety of biomedical implant settings.[20] Third, topographical growth cues are biologically relevant. Tissues present relevant microtopographies that growth cones must navigate. Both topographical and biochemical cues activate similar signaling to direct growth cone guidance behavior.[21,22] Finally, gradual, micron-scale, topographical cues can be engineered to offer similar or even greater influence on neurite guidance compared to biochemical cues.[15,16]

A significant question in neurite guidance by microtopographical features is what are the characteristics, abilities, and limits of a regenerating sensory neurite when turning in response to these cues.[23] This concept have been studied extensively, however, most commonly using diffusible biochemical cues, which lack the precision and reproducibility of engineered biophysical cues.[24,25] By using the precision and reproducibility enabled by topographically engineered cues, we can accurately define neurite turning behavior in response to a range of angles, as well as various amplitudes of these angled microfeatures. This reproducible system is essential for studying neurite turning since neurite outgrowth and growth cone migration are inherently stochastic with a wide variation of behavior across a population of neurons. Clarifying the consistency of how regenerating neurites can be effectively guided and determining what is the innate variation for this behavior are essential pursuits for informing the design future translational applications.[26]

For this work, primary sensory neurons were grown on novel topographically engineered substrates to present features with 6 different angle turn challenges. The amplitude of these features was varied by altering the photopolymerization parameters. To study neurite pathfinding generally, we assess neurite growth of both SGNs and DRGNs in response to this topography. The simple morphology of SGNs allows for detailed description of growth morphology, while the more dynamic DRGNs enable analysis of growth cone behavior and live imaging. In this work, we leverage the precision of photopolymerized topographical substrates to study the fundamental behaviors of neurite turning in response to a range of topographical cues, thereby informing the limits of pathfinding during neurite regeneration.

## 2. Methods

### 2.1. Micropatterned substrates

Topographically micropatterned substrates were generated as previously described using photopolymerization.[17] First, a monomer solution was formulated consisting of 40 wt% hexyl methacrylate (HMA, Aldrich), 59 wt% 1,6-hexanediol dimethacrylate (HDDMA, Aldrich), and 1 wt% of 2,2-dimethoxy-2-phenylacetophenone (DMPA, BASF). Then, to create the patterned substrates, this solution was evenly dispersed on a silane-coupled piece of cover glass by placing glass-chrome custom photomask (Applied Image Inc., Fig. 1b) on top. These samples were then exposed to 365nm light at an intensity of 16mW/cm^2^ using a high-pressure mercury vapor arc lamp (Omnicure S1500, Lumen Dynamics, Ontario, Canada) to polymerize the monomer solution (Fig. 1a). The opaque portions of the mask direct exposure of the UV radiation thereby modulating the rate of the polymerization locally to generate features on the surface. This process creates raised features or ridges underneath transparent bands where UV light intensity and the polymerization rate are highest. These substrates direct the growth of various cells and neurons, including SGNs and DRGNs.[27] Additionally, various geometries of topographical substrates can be generated by varying the photomask[24] or time of UV light exposure to change feature amplitude.[17] The duration of UV light exposure was varied in this work to create 3 microfeature amplitudes/depths of 2 µm, 4 µm, and 8 µm with the custom photomask.

**Figure 1:**
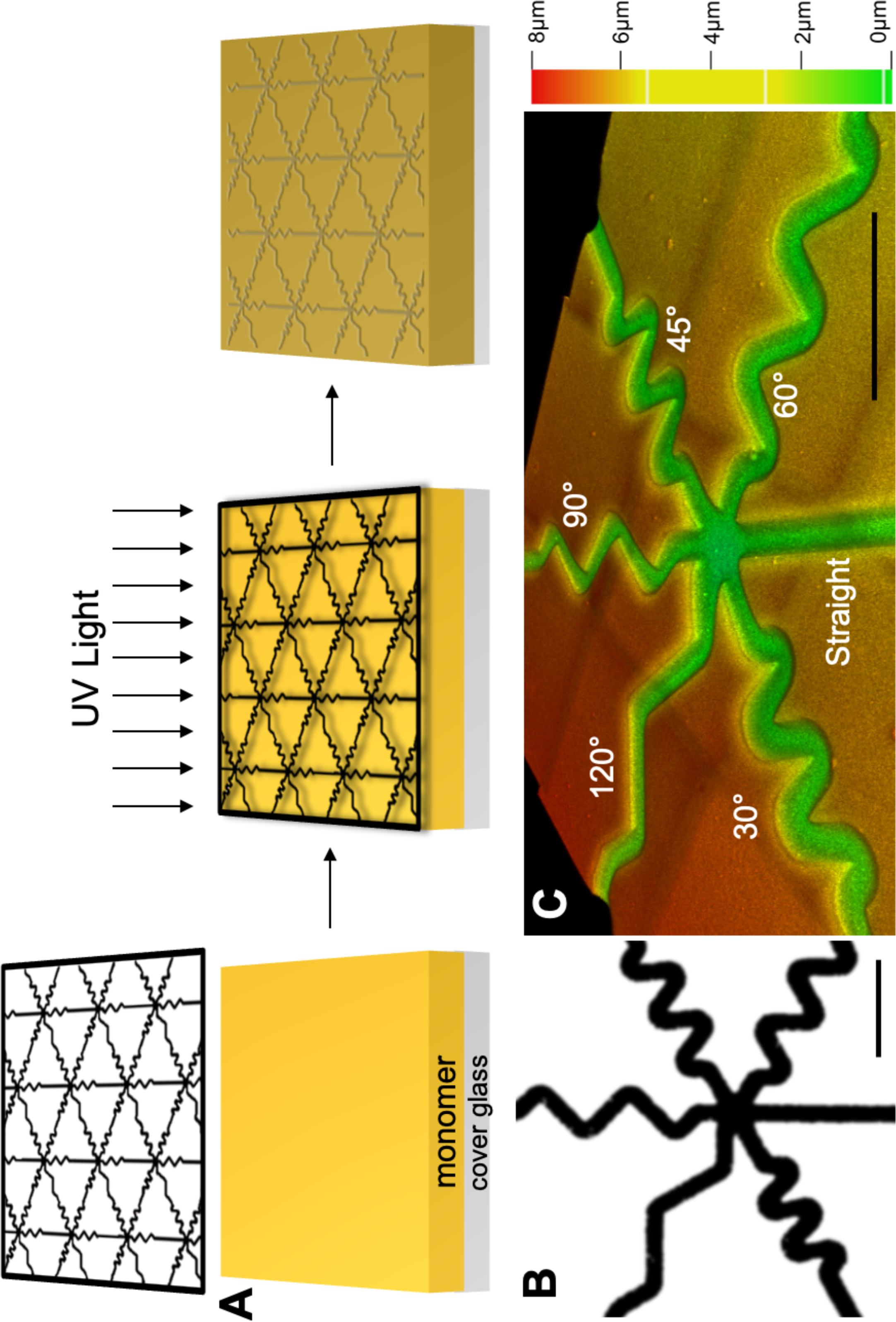
Schematic of Photopatterning Process and Pattern Characterization. A. Schematic of photopolymerizing micropatterned substrates. A monomer solution (yellow) is added to a silane coupled glass cover slip (grey). Then the photomask (black outlined pattern) is placed on top of the solution before exposing the system to UV light. The monomer polymerizes to form a solid substrate and the photomask is removed. B. The photomask (black). This pattern repeats over the photomask to create the topographically micropatterned substrate. C. A depth coded confocal microscopy image of multi-angled microfeatures patterned in the substrate surface. Scale bars = 100 μm.

To generate multiangle features and micropatterns, a novel photomask was used to generate the micropatterns utilized here (Fig. 1). This custom photomask consists of 6 different microfeatures, i.e., channels with gradually sloped edges, that are 20 µm in width arranged in a repeating hexagonal pattern. In addition to the control (straight) microchannel, the other 5 microfeatures consist of turns with varied angles (30°, 45°, 60°, 90°, and 120°) in which a neurite would need to deviate 20 µm perpendicular to the trajectory of the microfeature to navigate the angled turn (labeled in blue on Supp. Fig. 2a). For the microfeature nomenclature, we defined the microfeature in reference to the interior angle with 30° being the sharpest turn, 120° the most gradual, and 45°, 60°, and 90° representing intermediate turn challenges.

Prior work suggests that neurites can bypass the angle turn challenges in situations with zero or small overlapping distance (distance perpendicular to the direction of the microchannel in blue on Supp. Fig. 2a) in the zigzags.[24] Therefore, we chose to design each angle turn challenge to have an equivalent distance (20 µm) that the neurite would need to deviate perpendicular to the direction of the feature to navigate the turn. As such, the length of the straight portion of each microfeature’s zigzag is varied in order to standardize this design constraint.

### 2.2. Characterization of micropatterned substrate

The dimensions and characteristic of these micropatterned substrates were assessed using three methods. First, the topography of every substrate used in the following experiments was measured using white light interferometry (Dektak Wyko 1100 Optical Profiling System, Veeco; Supp. Fig. 1a). Feature amplitude was characterized in 5 regions of the glass coverslip and neurons were only cultured on areas within +/− 10% of the target microfeature amplitude. Vision software (Bruker) was used to create 3D images of substrates using this measurement technique.

A second method to characterize the micropatterned surface was scanning electron microscopy (SEM, Hitachi S-3400N; Supp. Fig. 1b). Conductive tape was used to mount sample substrates to SEM stubs and each sample was sputter coated with carbon prior to microscopy examination. The specimen was imaged top-down to assess the polymer surface.

The third method to assess the micropattern feature dimensions was confocal microscopy (STELLARIS 8, Leica; Fig. 1c, Supp. Fig. 1c)). This approach was possible since the polymer interacts with ultraviolet light, therefore, enabling it to be imaged via confocal microscopy using a 405nm laser. By creating a z-stack, the topographical dimensions as well as the character of the substrate surface can be determined.

### 2.3. Animals

All procedures involving animals were conducted in accordance with the NIH Guide for the Care and Use of Laboratory Animals and were approved by the University of Iowa Institutional Animal Care and Use Committee. All mice were maintained on a C57BL/6 (Envigo) background, housed in groups on a standard 12:12 h light: dark cycle with food and water provided ad libitum, and used at p3-5 of age, when SGN peripheral processes already contact hair cells in the organ of Corti. For live cell imaging, neurons were harvested from Pirt-GCaMP3 transgenic mice.[28]

### 2.4. Spiral Ganglion Neuron cultures

One day before culturing, substrates were sterilized by soaking in 70% ethanol for 2 min and then exposing them to UV light in a cell culture hood for 15 min. Cloning cylinders were placed on the patterned coverslip and then filled with poly-L-ornithine solution (Sigma-Aldrich) for 1 h at RT. After washing the substrate with sterile Milli-Q® water, a laminin solution was added (20μg/ml, Sigma-Aldrich) and incubated overnight at 4°C. SGN cultures were prepared from mice between post-natal day 3 and 5 (p3-p5). Mice were decapitated, and their cochleae were isolated from their temporal bones in ice-cold PBS. The spiral ganglia were then isolated and placed into ice-cold HBSS(-/-) as previously described.[29] Enzymatic dissociation was then performed in calcium- and magnesium-free HBSS with 0.1% collagenase and 0.125% trypsin at 37°C for 25 min. 100 µL of FBS was added to stop dissociation. Ganglia were washed with Neurobasal media before placing in supplemented Neurobasal Medium (Thermo Fisher) containing: 5% FBS, 2% N2 Supplement (Thermo Fisher), 10 µg/mL Insulin, 50 ng/mL BDNF (R&D Systems), and 50 ng/mL NT-3 (Sigma-Aldrich). Once in supplemented media, ganglia were triturated first with a 1000 µL pipette tip followed by a 200 µL tip. Cultures were plated onto micropatterned substrates and maintained in a humidified incubator with 6% CO_2_ for 48 h.

### 2.5. Replated Dorsal Root Neuron Cultures

DRGNs were isolated as previously described,[30] however, using neonatal (p3-p5) mice. First, a 24-well polystyrene plate was coated with poly-L-ornithine solution (Sigma-Aldrich) for 1 h at RT. The surface was washed with sterile Milli-Q® water three times prior to a laminin solution (20 μg/ml, Sigma-Aldrich) being added and incubated overnight at 4°C. After warming the coated well plate for 1 h at 37°C, the freshly dissected DRGNs were cultured on this plate for 72 h before being replated. The replating procedure has been described previously,[31] but after a 1 min incubation with TrypLE Express (Thermo Fisher), warm media was used to gently triturate the culture surface and lift the adhered neurons. The resulting replated DRGNs (rDRGNs) were then cultured on the micropatterned or unpatterned HDDMA/HMA substrates in a humidified incubator with 6% CO_2_ for 24 h.

### 2.6. Immunofluorescent labeling

After culture, media was removed and the culture surface was washed three times with phosphate-buffered saline without calcium and magnesium (PBS(-/-)). 4% paraformaldehyde in PBS(-/-) (Fisher Scientific) was added to fix the neurons. After 20 min at RT, cells were washed with PBS(-/-) three times, and then blocking buffer was added (5% normal goat serum [ThermoFisher], 0.2% Triton™ X-100 [Fisher Scientific], and 1% BSA [Research Products International] in PBS[-/-]). After incubating for 30 min at RT, different approaches were taken for the antibodies. For SGNs, mouse monoclonal anti-NF200 antibody (RRID: AB_260781, Millipore Sigma) was added (1:400 in blocking buffer) for 2 h at 37°C. Following this incubation, the culture surface was washed 3 times with PBS (-/-) and then Goat anti-mouse Alexa Fluor®546 (RRID: AB_2534089, ThermoFisher) in blocking buffer (1:800) was added for 1 h at RT in the dark.

For the rDRGNs, chicken polyclonal antibody to NF200 (RRID: AB_2313552, Aves) was added (1:800 in blocking buffer) for 1 h at RT. Following this incubation, the culture surface was washed three times with PBS (-/-) and then Goat anti-Chicken Alexa Fluor®546 (RRID: AB_2534097, ThermoFisher) in blocking buffer (1:1000) was added for 1 h at RT in the dark. Following the secondary antibody incubation, both systems were treated the same; cultures were again washed with PBS (-/-) three times. Lastly, the polymer-coated cover glass was mounted onto a glass slide using Fluoromount-G® (SouthernBiotech) and kept in the dark for 24 h before imaging.

### 2.7. Semi-automated measurement of neurite length within microfeature

Every SGN neurite that encountered a micropatterned, topographical microfeature was assessed in this analysis, in which neurite length in the microfeature was measured using NeuronJ.[32] Neurite length in microfeature was measured from where the neurite entered the microfeature to where it either exited the microfeature or the endpoint of the neurite was reached. These length data were then grouped both by turn angle and microfeature amplitude. For this approach, the unpatterned control condition was generated by overlaying the shape of the 60° microfeature over a flat portion of the substrate, and length data was measured in the same manner as neurites that encountered this pseudo-pattern.

### 2.8. Manual assessment of neurite pathfinding

Another method for assessing neurite pathfinding behavior was qualitatively scoring every neurite encounter with a microfeature. Neurites were scored as either turning or not turning. In particular, two conditions for which they were scored as not turning were: 1. did not follow the microfeature; neurite growth is not oriented in the direction of the pattern and the neurite crosses out of the microfeature nearly perpendicular to the microfeature’s direction, and 2. the neurite follows the straight portion of the microfeature but fails to complete the turn challenge and leaves the microfeature. Two categories of neurites were deemed to have successfully turned in response to the microfeatures: 1. the neurite follows the microfeature and navigates one turn; the neurite enters the microfeature more than 30 μm before the turn challenge and remains positioned in the microfeature 30 μm past the vertex of the turning challenge, or 2. the neurite navigates multiple turns and/or the shaft is reoriented where it is aligned across multiple of the zigzagging microfeatures. Like the previous metric, these data were grouped both by turn angle and microfeature amplitude.

### 2.9. Live DRGN imaging

Neurons growing in response to these turns were imaged live. To do this, rDRGNs were cultured from mice expressing Pirt-GCaMP3, a genetically encoded calcium indicator. These neurons were cultured on substrates with 4 μm and 8 μm amplitude microfeatures for 18 h. At that timepoint, neurites that were currently growing in a microfeature and actively engaging with the ridge of the feature were selected randomly and imaged every 15 s for 1.5 h to capture the growth dynamics. These videos were assessed by tracking the growth cones that were in the microfeature at the beginning of the recording and determining if they remained in the microfeature throughout the 1.5 h of imaging.

### 2.10. Comparing rDRGN growth cone morphology on micropatterned substrate

To study the morphology and behavior of neuron growth cones in response to topographical growth cues, replated DRGNs (rDRGNs) were cultured on a different topographically patterned substrate consisting of repeating rows of ridges and grooves (10 μm periodicity and 3 μm amplitude). These micropatterns offered a consistent pattern and thus a systematic methodology for this analysis. The substrates were made in the same manner as described previously.[17]

For this assessment, rDRGNs were cultured on these micropatterns and on unpatterned substrates, then immunofluorescently labeled as above; however, the secondary antibody step was modified to assess growth cone morphology. For the modified secondary antibody step, Goat anti-chicken Alexa Fluor®488 (RRID: AB_2534096, ThermoFisher) (1:800) and Alexa Fluor™ 546 Phalloidin (ThermoFisher) (1:200) in blocking buffer was added for 1 h at RT. Fixed growth cones were imaged via confocal microscopy (STELLARIS 8, Leica). Images of the growth cones were analyzed by two approaches using Imaris software. First, the prolaticity of the shape of the growth cone was computed as in Equation 1. For this analysis, the software approximates the shape of the growth cone as an ellipsoid and derives the prolaticity, which is a measure for how elongated the growth cone is. This shape was assessed since others have shown that the shape of the growth cone is influenced by substrate cues, thus the micropatterned substrate here may cause the shape to become elongated [33,34]

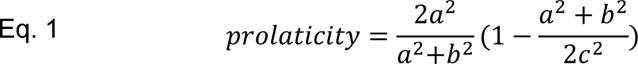

Second, the 2D shape of the growth cone was approximated as an ellipse and its major axis was derived. The angle made between this major axis and the neurite shaft was measured (Fig. 5).

### 2.11. Assessing SGN ability to remain in the microfeature around a turn

Since the neurons were encountering turn challenges of various microfeature amplitudes, the tendency of these neurons to keep their shaft in the microfeature around a turn was assessed. All the neurites that were deemed to have turned in the analysis above (section 2.8) were further studied and the proportion of these neurites that remained in the microfeature around the turn was determined.

For each microfeature turn angle, a mathematical definition of what characterizes a neurite holding its position around a turn was derived. For this definition, the greatest angle path a neurite could take from halfway down each straight segment and remain in the microfeatures was measured (Supp. Fig. 2b). This was found to be 134°, 138°, 142°, 151°, and 160° for the 30°, 45°, 60°, 90°, and 120° microfeature turn angles, respectively. ImageJ was used to measure the angle of the neurite’s shaft from the point on the shaft at the microfeature turn’s vertex to points on the shaft 30 μm on each side of the vertex (Supp. Fig. 2c). Neurites with angle of tension lower than that of the theoretical path of least tension for its given microfeature angle were characterized as holding tension, while those with angles greater were categorized as not holding tension. Neurite shafts that did not remain in the microfeature at the point of the vertex of the microfeature turn were also categorized as not holding tension. These data were grouped by microfeature amplitude.

In order to better visualize the neurites that held tension and/or reoriented, confocal microscopy was used to confirm the observations made by epifluorescence image analysis. The images were color coded by z-position to determine what plane the neurite shafts were positioned in these angled microfeatures.

### 2.12. Analysis of SGN neurite branching

Neurite branching was also assessed as a function of the angle at which the neurite encountered the edge of the microfeature. First, the angle at which the neurite encountered the ridge was measured using the angle tool in ImageJ (Supp. Fig. 5a,b). Then each neurite was scored as either follow, exit, or branch (if it branches at the microfeature boundary wherein and one branch follows and the other exits). For this approach, only the distal most neurite encounter with the micropattern for each neurite was assessed. Additionally, as part of this approach, the proportion of neurites that successfully turned was also measured as a function of the angle they approached the microfeature ridge and microfeature amplitude.

## 3. Results

### 3.1. Generation and characterization of topographically engineered substrate with multi-angled microfeatures

Using a customized photomask designed with varied angles (30° to 120°), photopolymerization techniques were used to generate methacrylate surfaces with the specified micropatterning consisting of grooves in masked regions (Fig. 1). By altering the duration of light exposure, patterns were engineered with microfeatures of varied amplitude, or depth (2 µm, 4 µm, and 8 µm). Thus, the photomask dictates pattern geometry while light exposure determines microfeature amplitude. The micropatterned substrates were characterized using multiple methods. The first method is via white light interferometry (Supp. Fig. 1a). This method enabled measurement of microfeature amplitude quickly and accurately. The substrates were also imaged with SEM to confirm that the microfeature ridges consist of sloping transitions rather than the typical, abrupt on/off features characteristic of other engineered micropatterns from lithography, etching, etc. (Supp. Fig. 1b). Lastly, z-stacks created with confocal microscopy also allowed for the measurement of the amplitude of the microfeatures, visualization of the smooth nature of the surface, and additionally the creation of depth color coded images to aid in the quantitative and qualitative assessment of the substrate (Fig. 1c and Supp. Fig. 1c).

### 3.2. Microfeature geometry determines the fidelity at which SGN neurites navigate microfeatures

The topographical multi-angled micropatterns enable the evaluation of neurite pathfinding in response to various turn challenges of precisely controlled geometry. In particular, SGN neurite pathfinding in response to 6 different microfeature turns (30°, 45°, 60°, 90°, 120°, and straight) was assessed across 3 different microfeature amplitudes (2 µm, 4 µm, and 8 µm). To provide an overall assessment of how strongly a neurite follows these varied micropatterns, we measured the distance that each neurite remained in the microfeature channel and compared the data with Two-way ANOVA. The analysis of neurite behavior on these substrates shows that the length of a neurite that remains in the microfeature increases both for more gradual turn challenges (straighter or larger angle microfeatures) as well as deeper microfeatures (Fig. 2). In particular, neurite length in microfeature increases an average of 20% from the 2 µm to 8 µm amplitude. Additionally, the length also increases by an average of 24% from the sharpest turn to most gradual (30° to 120°). Furthermore, we see similar findings when the neurite behavior at each type of turn challenge is scored as noted in section 2.8. The proportion of neurites that successfully navigate the turns increases with microfeature amplitude increasing when assessed with Two-way ANOVA. On average, the percent navigating the turns almost doubles from 18% making the turns on average in the 2 µm amplitude microfeatures up to 31% in the 8 µm amplitude microfeatures, a 72% increase in fidelity. In terms of microfeature angle, we see neurites are best able to turn in response to the moderate angle microfeatures (45°, 60°, 90°) (Fig. 3). This second analysis does not perfectly isolate the variable of turn angle since the straight portion of the microfeature varies in length for each turn. Thus, the conclusions of the Two-way ANOVA for this assessment are inconclusive and instead these data suggest that it is more challenging for a neurite to navigate a longer straight portion and more gradual turn (120° microfeature) than a shorter straight portion despite a sharper turn (45°, 60°, and 90° microfeatures). Lastly, we also see that regenerating neurites from another sensory neuron, rDRGNs, also successfully navigate these microfeatures in a geometry dependent manner (Supp. Fig. 3).

**Figure 2:**
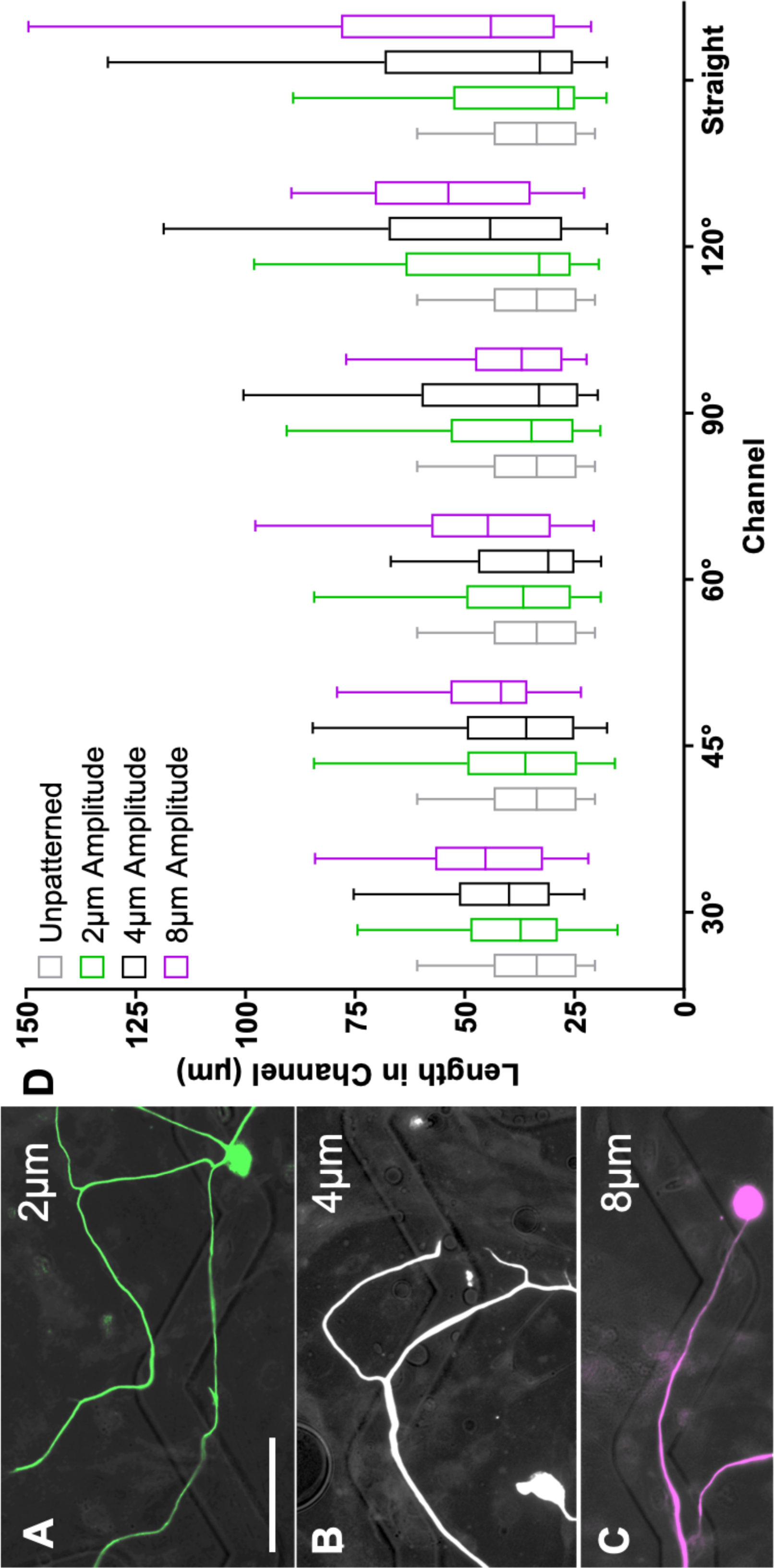
Feature Geometry Determines the Ability of SGNs to Navigate Complex Angle Microfeature Cues. A, B, C. Representative images of SGN neurite encountering a 120° angle microfeatures across the amplitude conditions A. 2 μm (green), B. 4 μm (white), and C. 8 μm (purple). D. 95% confidence interval of the distance that SGN neurites follow a microfeature once encountering that feature. n for each sub-condition ranges from 30 to 123. Unpatterned represents the shape of an angled microfeature superimposed onto a flat substrate. Two-way ANOVA shows that both microfeature amplitude and angle of turn affect SGN length in microfeatures with length increasing with deeper microfeatures and lower magnitude turns. p < 0.01. Scale bar = 50 μm.

**Figure 3:**
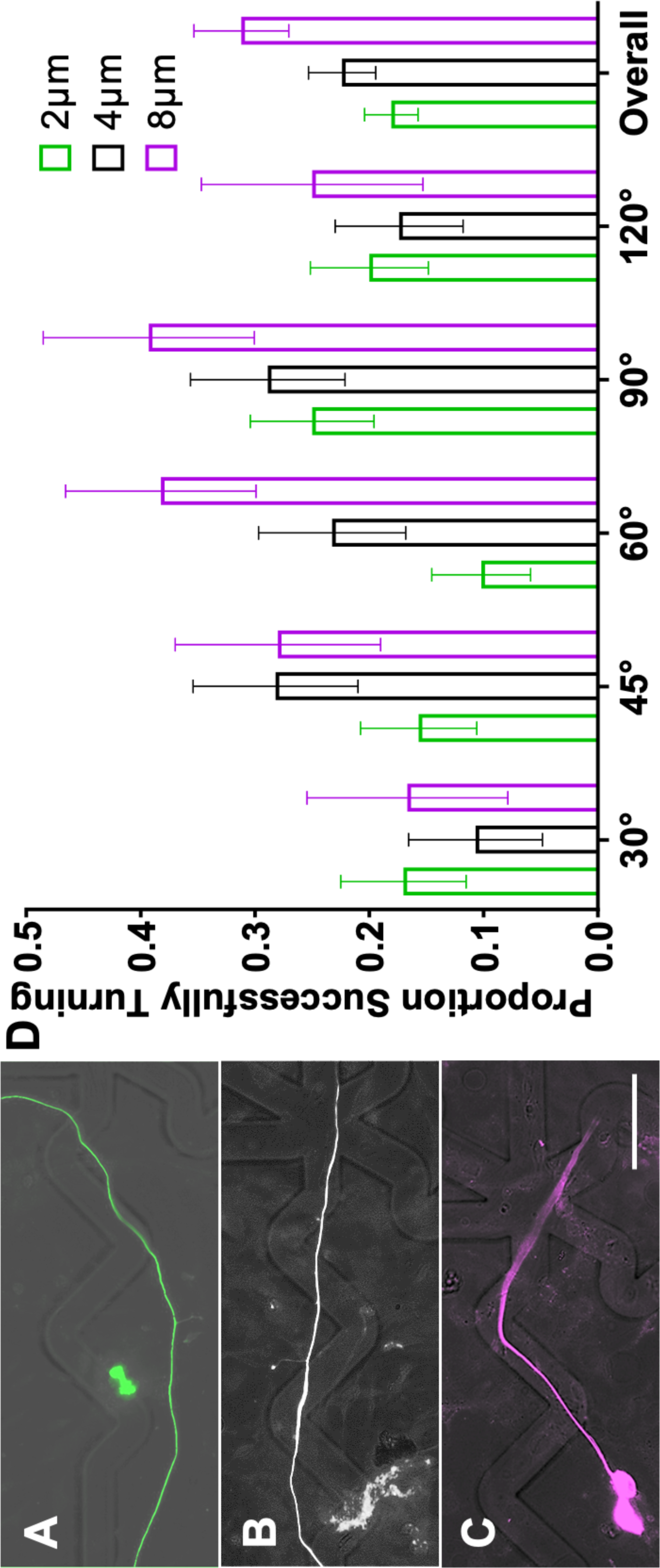
Microfeature Amplitude Promotes the Ability of SGN Neurites to Turn. A, B, C. Representative images of SGN neurite encountering a 90° angle microfeature across the amplitude conditions A. 2 μm (green) failing to turn, B. 4 μm (white) aligning across the microfeature, and C. 8μm (purple) making a turn. D. Proportion of SGN neurites that successfully navigate a turn in a microfeature when they encountered. n for each sub-condition ranges from 20 to 64. Two-way ANOVA shows that increasing microfeature amplitude improves the ability of SGN neurites to navigate turns. Error bars represent +/− SEM. p < 0.05. Scale bar = 50 μm.

### 3.3. Growth cones drive rDRGN neurite pathfinding

To better understand how neurites grow in response to these cues, replated DRGNs with a genetically encoded fluorescent indicator, Pirt-GCaMP3, were imaged live as they extended their neurites in the patterned angled microfeatures. In the videos, the growth cones were highly dynamic and strongly responded to the microfeature ridges in the 8 μm amplitude condition (Supp. Vid. 1). Based on the high fidelity with which the growth cones appeared to navigate the 8 μm amplitude microfeatures, we compared the proportion of the growth cones that remained in the microfeature throughout the duration of the 1.5h videos in the 4μm and 8μm amplitude microfeatures. Due to the low throughput of this analysis and the requirement that the growth cone of interest be actively engaging with a microfeature ridge at the beginning of recording, the growth cones analyzed were drawn from a random sample of turn angles and the growth cone position in that microfeature (Supp. Table 1). The proportion of neurites that remained in the microfeature was much greater in the 8μm amplitude microfeatures (95% +/− 5%) compared to the 4 μm (55% +/− 15%) (Fig. 4).

**Figure 4:**
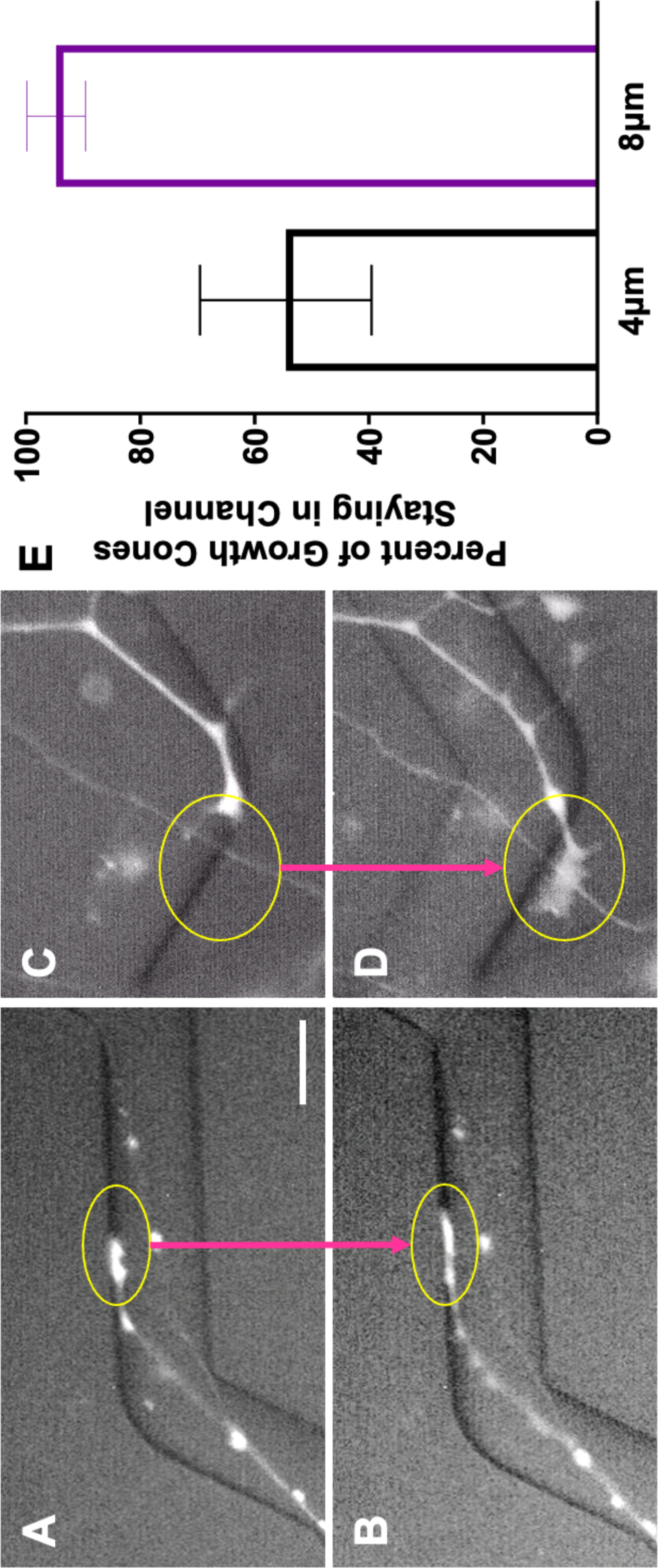
Growth Cones More Faithfully Remain in Deeper Microfeatures. A, B. Representative images from Supplemental Video 1a in which a rDRGN growth cone encounters and remains in an 8 μm amplitude, 120° angle microfeature. C, D. Representative images from Supplemental Video 1b in which a rDRGN growth cone encounters and exits a 4 μm amplitude, 120° angle microfeature. E. Percent of rDRGN growth cones that remained in the microfeature over the duration of 1.5 h recording. Z-test shows greater proportion remain in the microfeature in the 8 μm amplitude condition. Error bars represent +/− SEM. p < 0.01. Scale bar = 20 μm.

### 3.4. rDRGN growth cones on micropatterned substrates have distinct morphology

Given the stark difference in rDRGN growth cone pathfinding fidelity with different microfeature amplitudes, we further assessed growth cone behavior on micropatterned substrates by growing rDRGNs on a topographical substrate consisting of repeating rows of ridges and grooves (10 μm periodicity and 3 μm amplitude). We sought to determine if and how the growth cone morphology and orientation are altered when grown on topographically patterned substrates to inform how the strong differences in pathfinding in Fig. 4 manifest. When cultured on this micropatterned substrate, the ellipse shape of the rDRGN growth cones become more elongated and prolate than on unpatterned substrates, 0.431 (+/− 0.037) and 0.274 (+/− 0.043), respectively (Fig. 5c). Additionally, the angle difference between the major axis of the growth cone and the neurite shaft was smaller in the neurons grown on the micropatterned substrate, a median difference of 5° compared to 20° on the unpatterned substrate (Fig. 5d).

**Figure 5:**
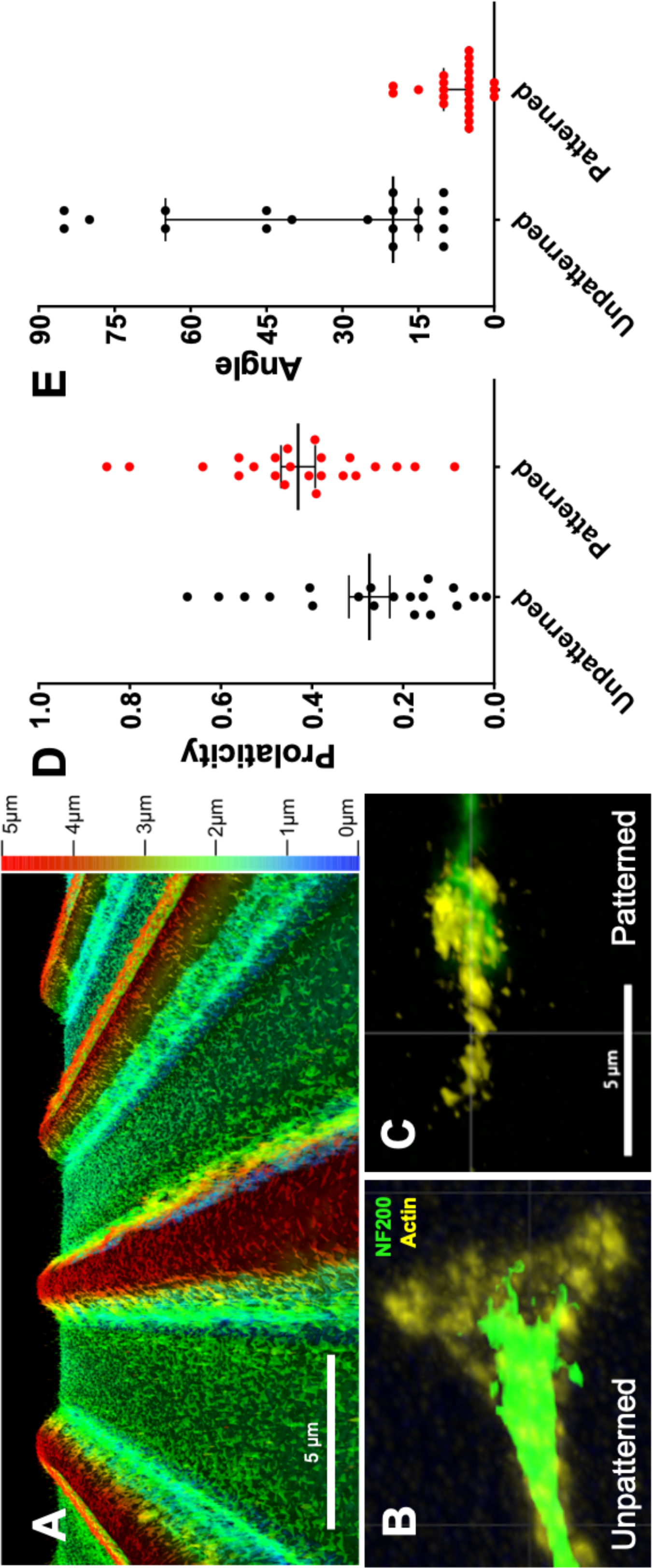
Growth Cones Exhibit Distinct Morphology and Behavior on Topographically Micropatterned Substrate. A. Depth coded confocal image of repeating rows of ridges and grooves substrate used for this experiment (3 μm amplitude and 10 μm periodicity). B. Representative image of a rDRGN growth cone grown on an unpatterned substrate. C. Representative image of rDRGN growth cone grown on the micropatterned substrate in A. D. Growth cone shape was approximated as a spheroid and its prolaticity was calculated as per Equation 1. T-test shows growth cones on the patterned substrate were more prolate. Error bars represent +/-SEM, p < 0.05. E. The major axis of this spheroid was found, and the angle difference between this axis and the neurite shaft was measured. Mann-Whitney shows that this angle was smaller for the neurons on the patterned substrate. Error bars represent 95% confidence interval. p < 0.001. Scale bars = 5 μm.

### 3.5. DRGN neurites reorient their shafts as they pathfind in response to microfeature turns, but not in a feature amplitude dependent manner

A significant subset of neurons aligned across, as opposed to with, the zig-zagged microfeatures in the fixed sample studies (Supp. Fig. 4). To dynamically assess the behavior leading to this observation, live imaging of neurite shafts was conducted. The live imaging demonstrates that the neurite shafts are constantly reorienting their position as the neurite is elongating. The neurite shafts appear to be mobile and drift as the growth cone pathfinds. In particular, the neurites tend to reposition their shaft such that it is becomes aligned in the direction of elongation. This movement thereby minimizes the number of neurites that hold their position in a zigzag or hold their tension around a turn. This behavior occurs since the neurite shaft moves to reposition itself outside of the microfeature and becomes more aligned in the direction of neurite elongation (Supp. Vid. 2). Additionally, via SEM of these rDRGNs display neurites tightly following the microfeature ridge or just hopping over the vertex of the angle challenge of the microfeature (Supp. Fig. 4).

The neurons that successfully turned in Fig. 3 were again assessed to determine if the proportion of neurites reorienting around the turns was related to microfeature geometry. In this analysis, there is no significant effect of feature amplitude on neurite shafts remaining in the microfeatures when making turns. Though a higher proportion of neurites remain in the microfeature when turning in the 8 μm amplitude condition, this difference is not statistically significant by One-way ANOVA since we are comparing small subsets of the data and the magnitude of the difference is not large (31% vs. 46%, respectively) (Fig. 6). Additionally, no significant relationship was found between angle of the turn and the ability of a neurite to hold tension when making turns.

**Figure 6:**
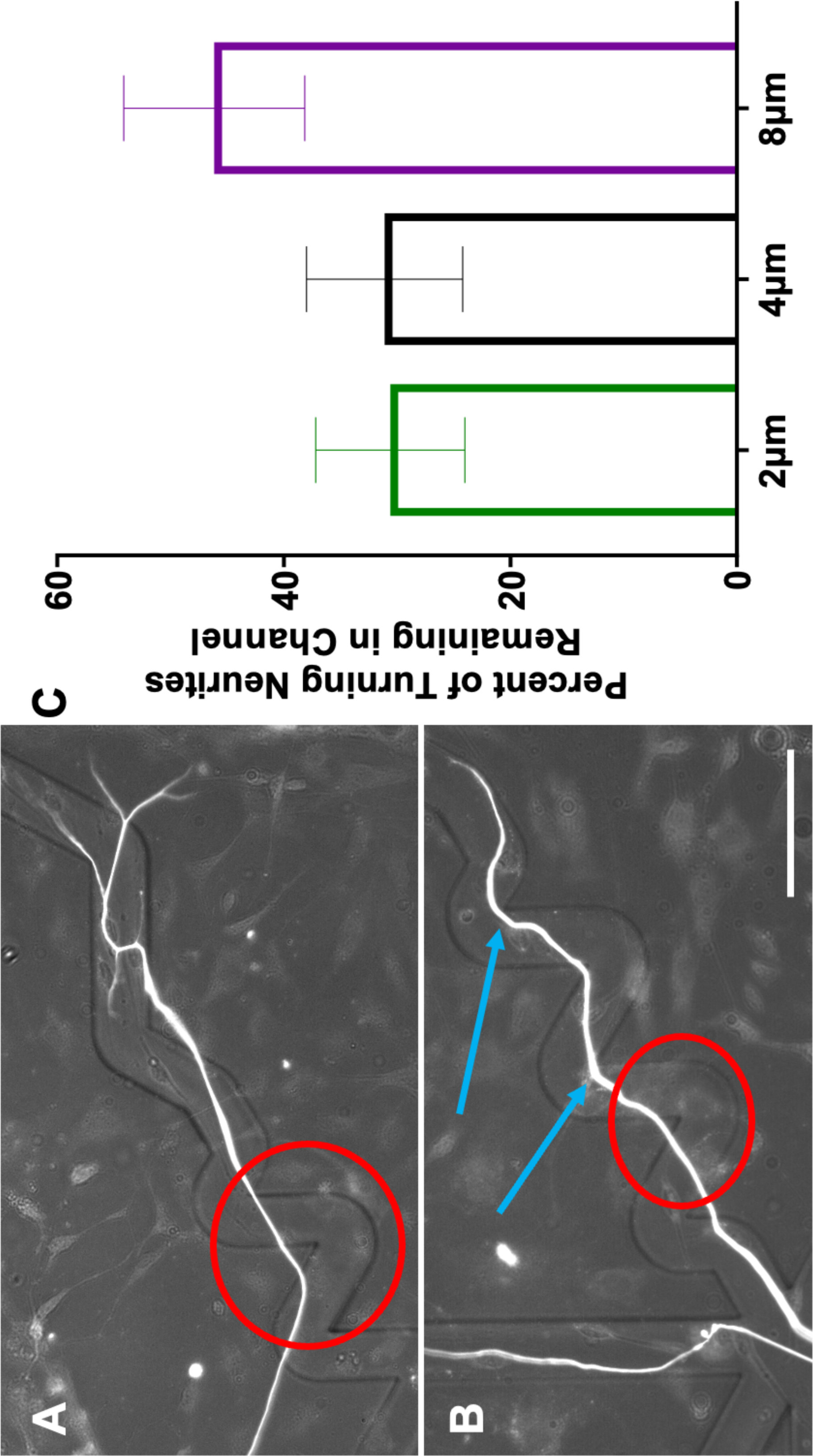
Neurite Shafts Remain in Microfeatures at Similar Rates when Navigating Turns Regardless of Feature Amplitude. A. Representative image of DRGN neurite unable to hold position around a turn through a 4 μm amplitude, 60° angle microfeature. B. Representative image of DRGN neurite holding position around 2 turns through an 8 μm amplitude, 60° angle microfeature. Red circles indicate neurite shafts not holding the turn while blue arrows show neurites that do hold the turn. C. Percent of neurites that successfully navigate the angle turn challenge and with shafts holding that position. One-way ANOVA shows no difference in treatment groups (p = 0.24). n = 49, 45, and 39. Error bars represent +/− SEM. Scale bar = 50 μm.

### 3.6. Neurite branching is related to turning efficacy

During live imaging, neurites were observed to branch when the growth cone encountered a ridge or turn challenges on the angled microfeatures. Additionally, the growth cones were observed to have a branched, arborizing appearance as they navigated the microfeatures (Supp. Vid. 2). To quantitatively assess neurite branching, we scored neurite behavior (follow, exit, or branch) as a function of microfeature amplitude and the angle at which the neurite encountered a topographical ridge from within the angled microfeatures (Supp. Fig. 5a,b). This modified approach was used since the exact angle at which a growth cone encounters the feature ridge varies within each microfeature condition. Firstly, the results of this modified quantification echo the findings of Fig. 2 and 3 showing that the proportion turning increases with greater microfeature amplitude and with more obtuse angles (Supp. Fig. 5c). The neurites are observed to better follow the microfeatures by a magnitude of at least 3.5-fold when they approach the microfeature ridge at a less severe angle. Additionally, neurites are seen to better follow as microfeature amplitude increases from 2 µm to 8 µm by an average of 2.8-times (Supp. Fig. 5c).

When studying the relationship between neurite turning and branching using this modified assessment, we see neurite branching occurs most frequently when the chance of neurite turning approximated 50% (Fig. 7). Branching is observed to be less likely to occur when the neurite turning outcome becomes more certain, i.e., when approaching no chance of turning or all neurites making the turn. The microfeature angle where branching was maximized varied across microfeature amplitudes. For the 2 µm microfeatures, maximal branching occurs in gradual turns (15° - 30°), 4 µm in moderate turns (30° - 45°), while sharper turn challenges maximized branching in the 8 µm amplitude microfeature (75° - 90°).

**Figure 7:**
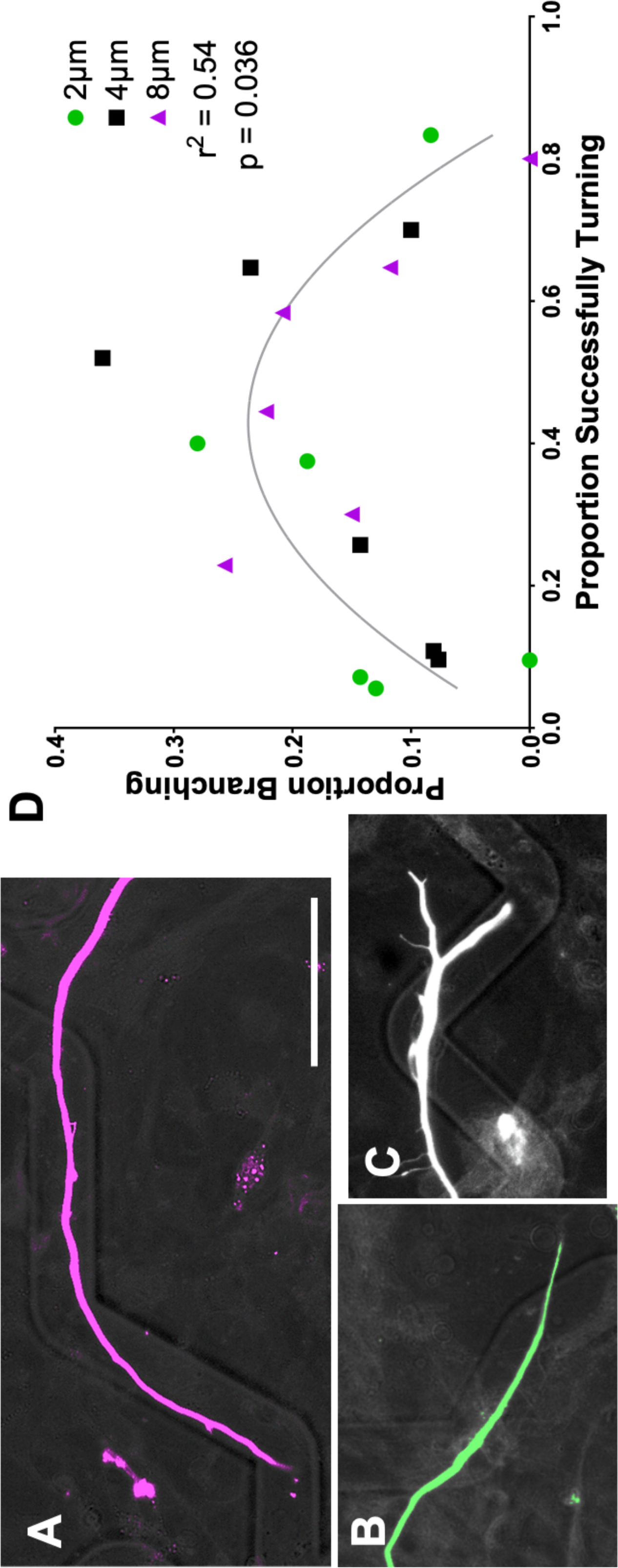
Neurite Branching is Most Likely When Neurite Turning Result is Uncertain. A. SGN neurite turning in response to an 8 μm amplitude, 120° angle microfeature, representing an “easy turn” toward the far right of the graph. B. SGN neurite exiting the microfeature in a 2 μm amplitude, represents a “challenging turn” towards the left of the graph. C. SGN neurite branching to both follow and exit the microfeature in a 4 μm amplitude, 60° angle feature, representing an intermediate challenge where branching was seen to be most likely. D. Neurite branching plotted as a function of neurites successfully turning. A second order polynomial fits the data with r^2^ = 0.54 which equates to p = 0.036.

## 4. Discussion

Neural prostheses including CIs can restore lost function, however, the auditory perception and quality provided is significantly lower resolution compared to native pathways. CIs represent the most successful neural prostheses to date, but the large distance between the stimulating electrode and the target SGNs limits their function. Closing this physical gap to improve signal resolution is a primary goal with a variety of approaches proposed including; (1) altering the electrode array shape to place it in a perimodiolar position so that the stimulating electrodes reside closer to the neurons,[35,36] and (2) tissue engineering endeavors to induce de novo, organized neurite growth towards the stimulating electrodes.[11,37] For optimal translational potential of the latter approach, shelf stability, biocompatibility, and precision of guidance should be prioritized.[38] Thus, in this work we use the spatial and temporal control inherent to photopolymerization to create novel, precise topographically patterned substrates to guide neurite regeneration and thereby evaluate the ability of sensory neurons to pathfind in response to topographical angled turn challenges for both SGNs and rDRGNs to use diverse neurons with different characteristics.

Herein, we have demonstrated a novel system to assess the ability of neurites to turn in response to topographical cues of varied angles and amplitudes (depths). Using engineered micropatterned substrates, we see that the geometry of the angle turn challenge, i.e., microfeature amplitude and angle of turn, determines the ability of the sensory neurites to navigate the turn and follow the micropattern (Figs. 2 and 3). This conclusion appears straightforward; however, we see a wide variation in how neurites respond to the same geometric cue (Fig. 2). Furthermore, neurites can follow or hop out of any microfeature studied (Fig. 3) implying that in these microfeature turns there is not a clear ceiling or floor of the ability of a neurite to navigate a turn. Therefore, while the geometry of the angled features influences how well sensory neurites track the microfeature guidance cues (Fig. 2) and navigate the turns (Fig. 3), the process by which these neurites navigate these turns is not simply deterministic. These data suggest that as SGN neurites navigate complex turn challenges, their behavior reflects a balance of deterministic cues guiding their growth and their innate stochastic outgrowth patterns. Therefore, the strength of the guidance cue does not determine growth cone behavior, but rather shifts the probability distribution of growth cone turning behavior.

This stochastic nature of growth cone motility has been described previously.[39,40,41] In such foundational work, neurite pathfinding results from an interplay between the strength of the turn cue,[15,17] stochastic actin polymerization,[41] and signaling from the support cells from the primary culture influencing guidance cues.[42] Within the experimental system, these factors result in non-uniform pathfinding observations within each geometric condition, e.g., neurites do not navigate the gradual turns in the 8 μm amplitude with perfect fidelity. It is important to note, that sufficiently strong cues can overpower the stochastic nature of the neurite pathfinding to create nearly deterministic systems.[17,43] While the data here do not show absolute determinism, we show that topographical cues, of biologically relevant dimensions, can strongly shift the probability distribution of this system. Thus, this is an appealing model to study turning and guidance since it reflects the balance of determinism and stochasticity inherent to neurite guidance *in vivo*.[44,45] Such a model of neurite turning in response to topography can be applied to study fundamental signaling mechanisms that contribute to neurite guidance and translation to *in vivo* applications.

To better understand the neurite pathfinding process, live imaging of rDRGN neurites navigating the microfeatures was conducted (Supp. Vid. 1,2). In these videos, the growth cones had a strong tendency to faithfully track in the microfeature for the 8 μm amplitude microfeatures (95%), while in the shallower 4 μm microfeature, the fidelity to the microfeature was less consistent (55%) (Fig. 4). These data suggest two main findings. First, the angled microfeature system can be engineered that is much more deterministic than stochastic in guiding neurite growth. Growth cones consistently remain in deep microfeature (8 μm amplitude). Thus, strong, biologically relevant topographical cues can be engineered that precisely direct neurite guidance and turning. Second, based on the significant difference observed in Fig. 4., it appears that growth cone behavior accounts for the differences in pathfinding that is observed in the prior analyses, Figs. 2 & 3 and Supp. Fig. 3.

Thus, after observing differential rDRGN growth cone behavior with different amplitudes of the various angled microfeatures, we further assessed how growth cone morphology and behavior change using a topographical substrate with narrow repeating rows of ridges and grooves. This system offered a more systematic methodology and further support for growth cone behavior being involved in differential pathfinding to topographical cues, since the growth cone is more prolate when grown in the microfeatures (Fig. 5c). Furthermore, we see that the growth cone is elongated to be strongly oriented in the direction of the microfeatures and neurite shaft (Fig. 5d). The alignment of the growth cone and shaft suggests that changes in growth cone morphology and behavior underlie the pathfinding observations downstream. Additionally, by integrating these findings with previous work, it appears that the patterned substrate and growth cone interact to cause the growth cone to become elongated in the direction of the microfeatures. This prolate shape, drives growth in the direction of the major axis of the ellipsoid growth cone, which is in line with the ridges and grooves. The neurite shaft then orients in the same trajectory since the growth cone is uniformly driving growth in that direction.

It is evident that topographical features change rDRGN growth cone behavior and that the growth cones have differential responses across topographical feature amplitudes. It is unclear if the behavior of the growth cone is the main driver of SGNs differentially navigating the various amplitude microfeatures since we are unable to directly assess their growth cones and other observations may complicate this conclusion. One complication is that the neurite shafts appear tremendously dynamic and reorient around the turns as the growth cone traverses the environment when imaged live (Supp. Vid. 2). The neurite shafts reorienting their position may obscure the inferences of our work since; 1) how a shaft is oriented at a given timepoint may not be representative of the path a growth cone took to create that neurite and 2) the neurite shaft may remain in the microfeature around a turn at a slightly higher rate in greater amplitude microfeatures (Fig. 6). In combination, these two points bring the findings of Figs. 2 and 3 under scrutiny. First, it is important to note that even with this chaotic reorientation occurring, we still see clear and significant differences in the corresponding data. The clear results in Figs. 2,3 and Supp. Fig. 5 strengthen the conclusion that microfeature geometry determines neurite turning since even with reorientation convoluting the neurite tracing data, the effect is still clear. However, the question of what is the main driver of the observed effect remains, the differential behavior of the shafts or the growth cones. While neurite shafts may reorient less frequently in the 8 μm amplitude microfeatures, no significant difference is observed between the 2 μm and 4 μm (Fig. 6). Furthermore, the magnitude of the effect observed in the shaft reorientation measure (Fig. 6) is much smaller than the growth cone assessment (Fig. 4). Comparing these data implies that the differential behavior of the growth cone likely outweighs the effect of neurite shaft reorientation in driving the observed pathfinding differences. That being said, the reorientation of the neurite shafts is an important finding that motivates a reinterpretation of prior studies using fixed neurons to study pathfinding in response to various cues.[24,46] Our data suggest that the growth cone may be consistently navigating a microfeature, but the shaft may reorient bringing it out of the microfeature. Therefore, tracing fixed neurites may not represent the true trajectory that the growth cone took in its elongation.

Beyond influencing neurite pathfinding, we also evaluated the relationship between the tendency of a neurite to branch and the ability of neurites to navigate a turn on these micropatterned surfaces.[47,48] We observed that branching occurred most often when the probability of neurites successfully making a given turn was essentially a tossup, near 50% (Fig. 7). When interpreted in combination with the live imaging data, these data suggest that the dynamic growth cones initiate branching at decision points on the micropatterns. When the two paths offer equal resistance, i.e. making a turn or exiting out of the microfeature, both neurite branches are more likely to persist.[49] By contrast, on micropatterns with stronger guidance cues and greater certainty of growth behavior, the neurite either never initiates a branch, e.g., the growth cone remains in the microfeature for a gradual turn in a high amplitude feature, or ends up retracting the branch in the less favorable path, e.g., growth cone easily hops out a shallow microfeature at a sharp turn.

Though our work does not directly address the causes of branching is occurring, it complements other work informing the process. First, others have indicated mechanical force plays a significant role in neurite branching, in that neurons retract a growth cone branch if sufficient mechanical stress is applied.[50] In this context, mechanical stress on the growth cone from either the topographical microfeature ridge or turn likely initiates growth cone branching or dictates which arbor to retracts. Additionally, topography has been implicated in branching frequency where topographical cues were presented to promote neurite branching.[51] The ability to promote branching is consistent with the findings here that branching does not occur randomly, but can be stimulated to an extent by precise cues presenting a challenge with two equally likely directions of growth simultaneously. Overall, this angled microfeature system, or similarly constructed cues, offer a useful model to study neurite branching decisions.

This work is one of the first to use an engineered topographically patterned substrate like this (Fig. 1), thus there are a few limitations of this initial work. First, neurons used in this study are derived from primary cell cultures. This means that the cells on the micropatterned substrate are a heterogeneous mixture of the various cell types present in the ganglia. Therefore, the topographical growth cues inherent to the substrate are not the only guidance cues the neurites encounter in this system, as the glial cells may affect neurite guidance as well.[42] This tradeoff is necessary in order to study the pathfinding behavior of primary neurons with their full expression of receptors and channels. Second, all data are from neurons derived from neonatal animals. Neonatal SGNs are more dynamic compared to adult SGNs, the target cells of patients receiving CIs. This shortcoming is necessary as the yield and growth of SGNs from adult mice is limited. Likewise, growth cone size and dynamics are enhanced in neonatal DRGNs. Thus, using neonatal DRGNs for the live imaging and growth cone studies is necessary. That being said, adult SGNs and DRGNs have been shown to respond to topographical growth cues as well so the results are expected to translate.[27]

## 5. Conclusions

Understanding the ability of neurites to sense and turn in response to topographical cues is critical for informing the future design of neuroprostheses such as CIs, neural conduits, and mechanisms for guiding neurite outgrowth *in vivo*. In these translational applications, neurites will be guided in 3D as opposed to on 2.5D substrates in this work. That being said, this work demonstrates fundamental principles that will be essential for the design of 3D systems for use *in vivo,* as the microfeatures in our work have sloped edges, so when a growth cone is growing within a microfeature for all intents and purposes the growth mechanics are the same as within a closed 3D tube with the same curvature.[17]. First, based on the decreased ability of neurites to pathfind to sharper angles, turns should be engineered to be gradual and to minimize zig-zagging as neurite shafts appear to resist this reorientation. Second, since the growth cones change their morphology and migration behavior to drive pathfinding in response to these microfeatures, translational 3D biomaterials ought to have aligned through-pores, which have been proposed to be advantageous for neural conduits compared to a random meshwork.[52,53] Lastly, minimizing neurite branching at the electrode interface will be necessary to optimize signal resolution in next generation CI. Our work here implies that branching occurs as function of neurite growth uncertainty when presented with multiple paths of growth. Therefore, in future material designs, instructive growth guides should be of sufficient strength to minimize exuberant branching. This may be challenging to accomplish since branching may occur at positions with unclear cues, such as when entering and exiting the designed material. Thus, as with pathfinding, neurite branching must also be considered, and minimized, for optimal recapitulation of the architecture and function of native neural system where growth is being induced.

## 6. Acknowledgements

The Pirt-GCaMP3 transgenic mice were gifted from Dr. Xinzhong Dong of John Hopkins University.

## 8. Supporting Information

**Supplemental Figure 1: Three Methods Used to Characterize the Engineered Multi-Angled Microfeatures on the Substrate Surface.** A. Depth coded white light interferometry image taken using an optical profiling system. B. Depth coded confocal microscopy image of 60° angle microfeature. C. SEM image of 60° angle microfeature and substrate surface. Scale bars = 50 μm.

**Supplemental Figure 2: Dimensions of Interest in the Multi-Angled Microfeature Substrate.** A. SEM image of 60° angle microfeature showing the 20 µm (blue) that a growth cone needs to deviate to navigate that turn. B. SEM image of 90° angle microfeature showing the measurement of the minimum angle required for a neurite to hold tension. This minimum angle was defined as the path with the most gradual angle a neurite could take from halfway down each straight segment and remain in the microfeature around a turn (151° for the 90° microfeature shown here). C. NF200 labeled SGN making and holding a turn in a 90° microfeature. Labeled in orange is the measurement of the angle the neurite makes around the turn found by the angle made with 30 µm of neurite length on each side of the turn. Scale bar = 50 μm.

**Supplemental Figure 3: Replated DRGN Neurites Also Follow These Microfeatures in a Geometry Dependent Manner.** A. Representative image of rDRGN fixed on an 8 μm substrate. B. 95% confidence interval of distance rDRGN neurite follows a microfeature once encountering it. n for each sub-condition ranges from 60 to 128. Unpatterned represents the shape of an angled microfeature overlayed onto a flat substrate. One-way ANOVA shows that distance the neurite remains in a microfeature increases with more gradual turns. p < 0.001. Scale bar = 100 μm

**Supplemental Table 1: Description of behavior of all growth cones imaged and assessed for Figure 4.**

**Supplemental Figure 4: Depth Coded Confocal Images Show z-Position of Neurite Shafts in Turns.** Images represent depth coded signal of combination of fluorescent signal from NF200 (568nm wavelength) and HMA/HDDMA micropatterned substrate (405nm wavelength). A. Replated DRGN neurite which partially exits the microfeature during a turn in an 8 μm amplitude, 90° angle microfeature. B. Replated DRGN neurite holding position on the sloped edge of a microfeature ridge in an 8 μm amplitude, 60° angle microfeature. Scale bar = 20 μm

**Supplemental Figure 5: Neurite Turning is Dependent on Angle the Neurite Approaches Microfeature Wall and Microfeature Amplitude.** A. Representative image of a SGN neurite approaching the edge at ∼85° and exiting the microfeature. Angle is calculated from the average trajectory of the neurite approaching the edge (orange arrow) and the line tangent the edge (blue). B. Representative image of a SGN neurite approaching the at ∼65° and turning. C. SGN neurite turning assessed as a function of the angle at which the neurite encounters microfeature wall, regardless of the angle of the microfeature in which the neurite was pathfinding. Two-way ANOVA shows that proportion of neurites increases with smaller angle needed to navigate the turn and with greater feature amplitude. p < 0.001

**Supplemental Videos 1: DRGN Growth Can Remain in (or Hop Out of) the Microfeature in Response to Turn Challenges.** A. Replated DRGN growth cone encountering and remaining in an 8 μm amplitude, 120° angle microfeature. B. Replated DRGN growth cone encountering and exiting a 4 μm amplitude, 90° angle microfeatures. Scale bar = 20 μm

**Supplemental Videos 2: Replated DRGN Neurites Reorient Their Shafts Across the Turns while Pathfinding.** A. Replated DRGN reorienting its neurite shaft across a turn in an 8 μm amplitude, 60° angle microfeature. B. Replated DRGN reorienting its neurite shaft across a turn in a 4 μm amplitude, 60° angle microfeature. Scale bar = 20 μm

