## Supplementary figures and images for "The geometry of photopolymerized topography influences neurite pathfinding by directing growth cone morphology and migration"

### Supplemental Figure 1

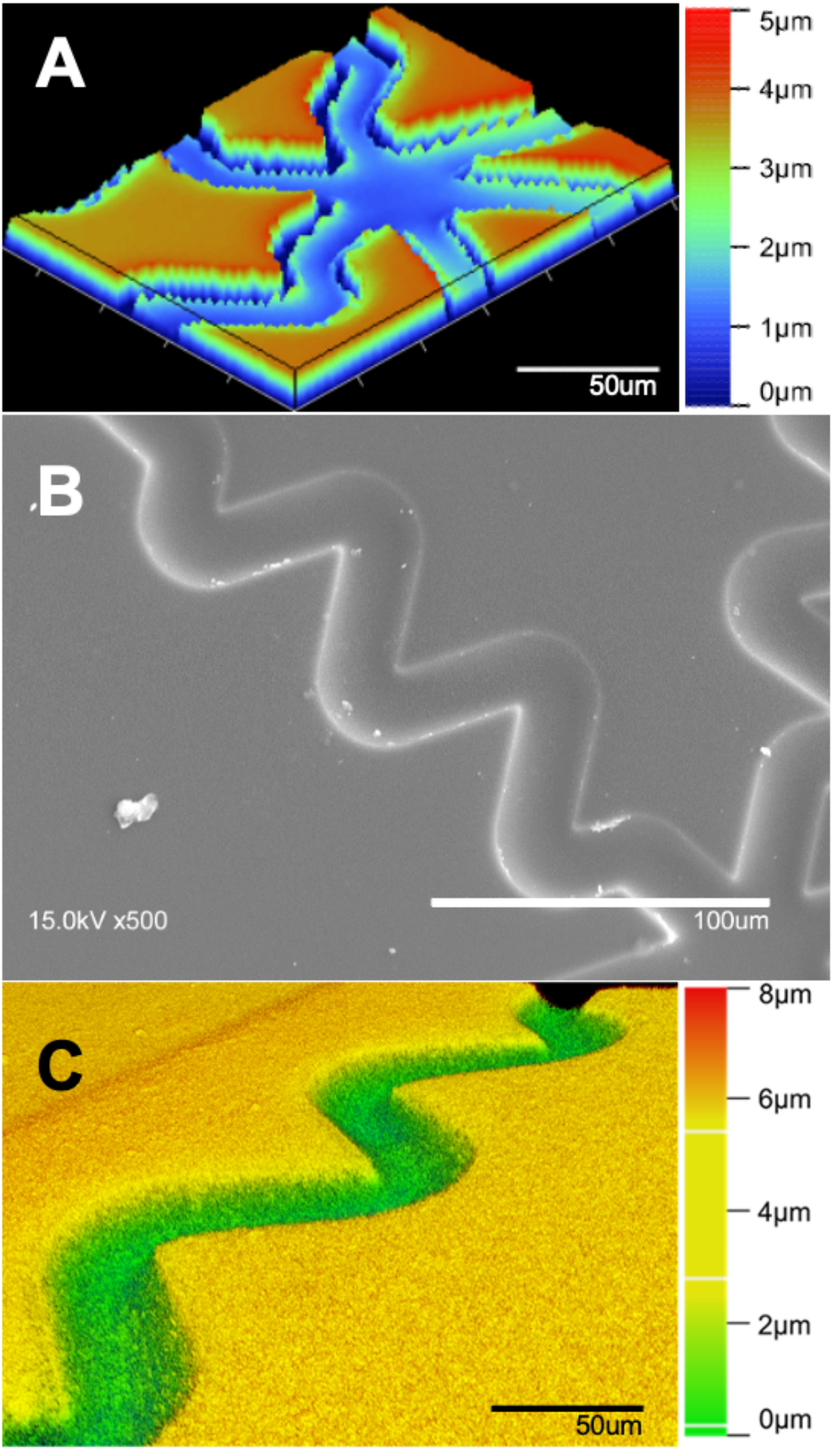

### Supplemental Figure 2

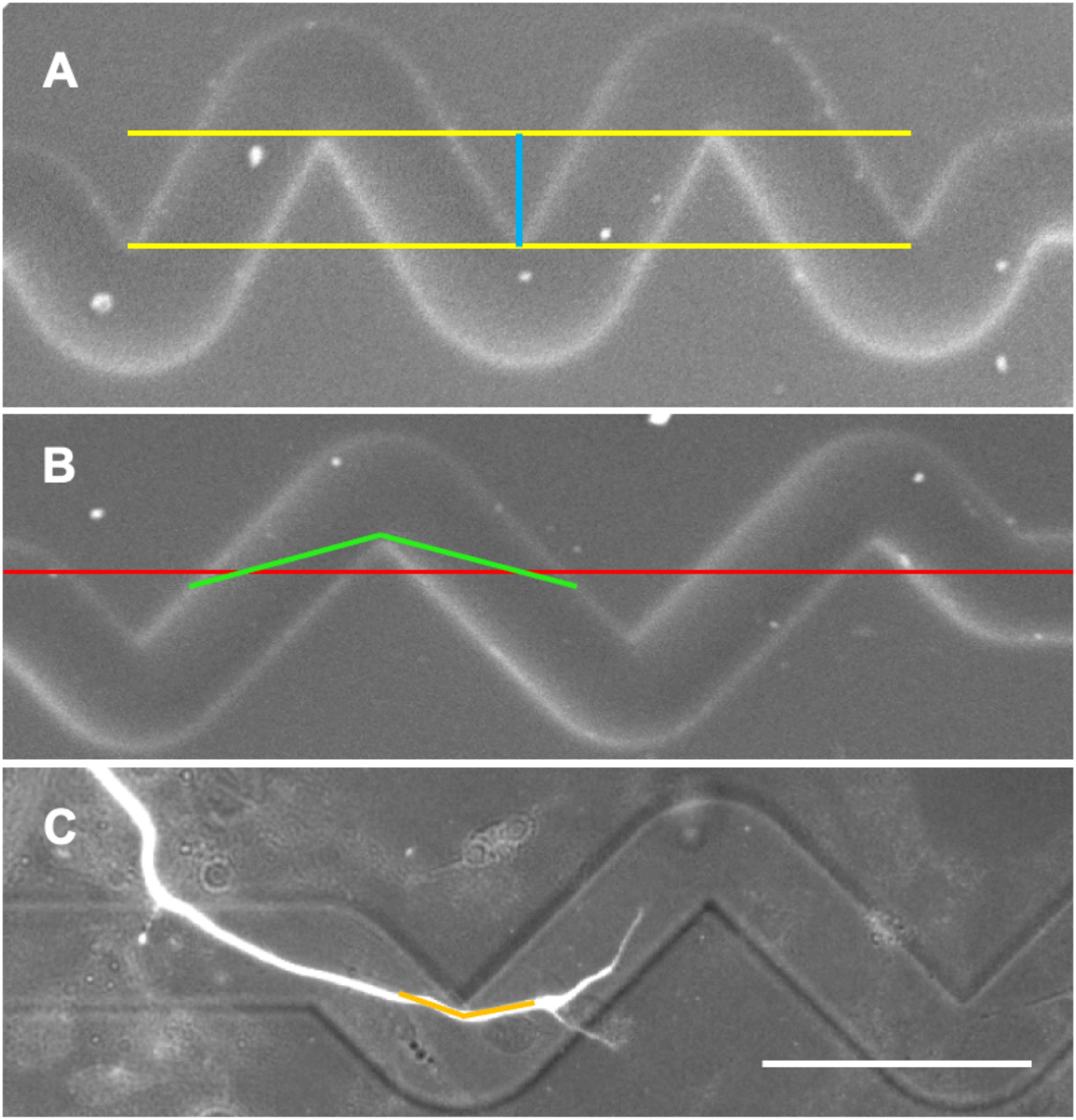

### Supplemental Figure 3

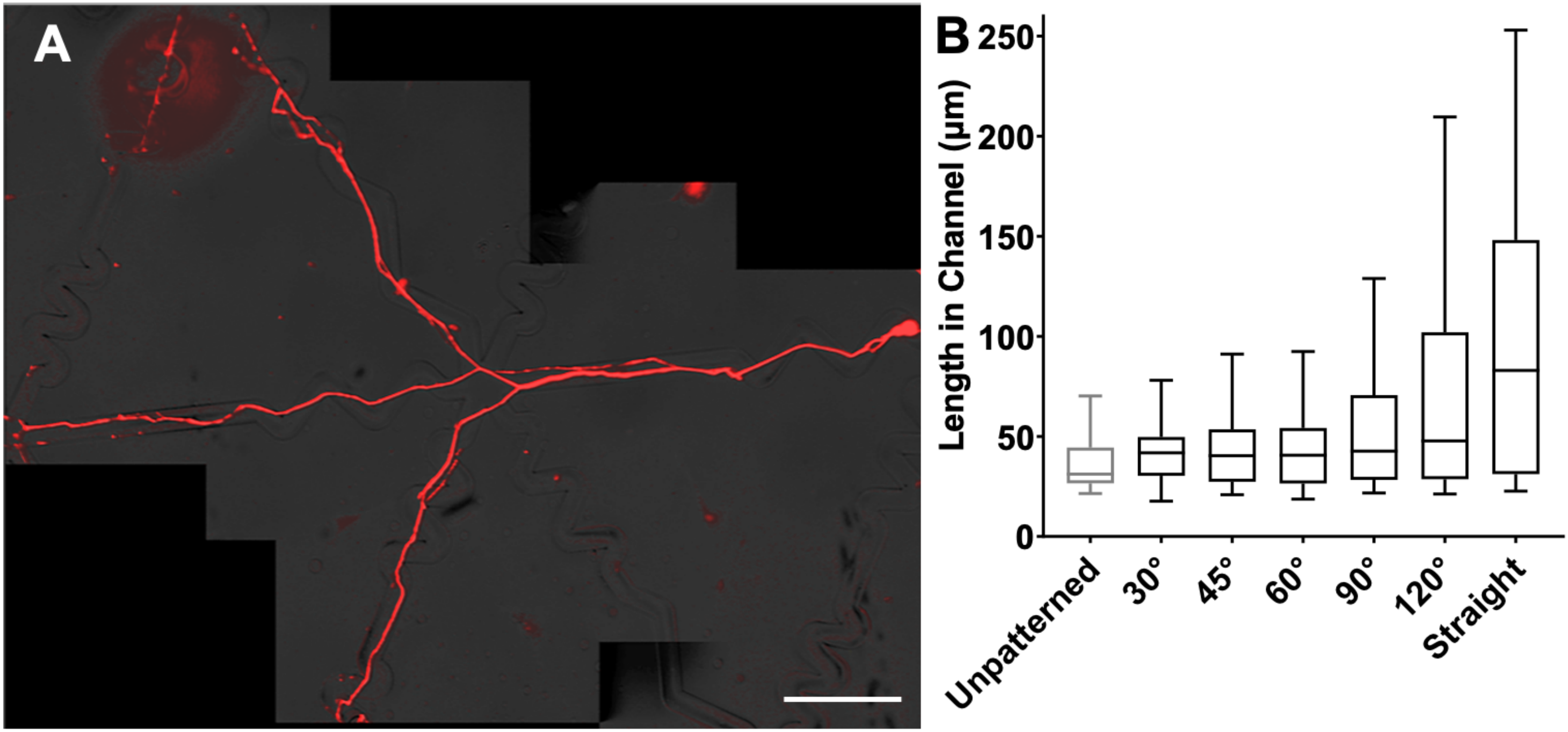

### Supplemental Figure 4

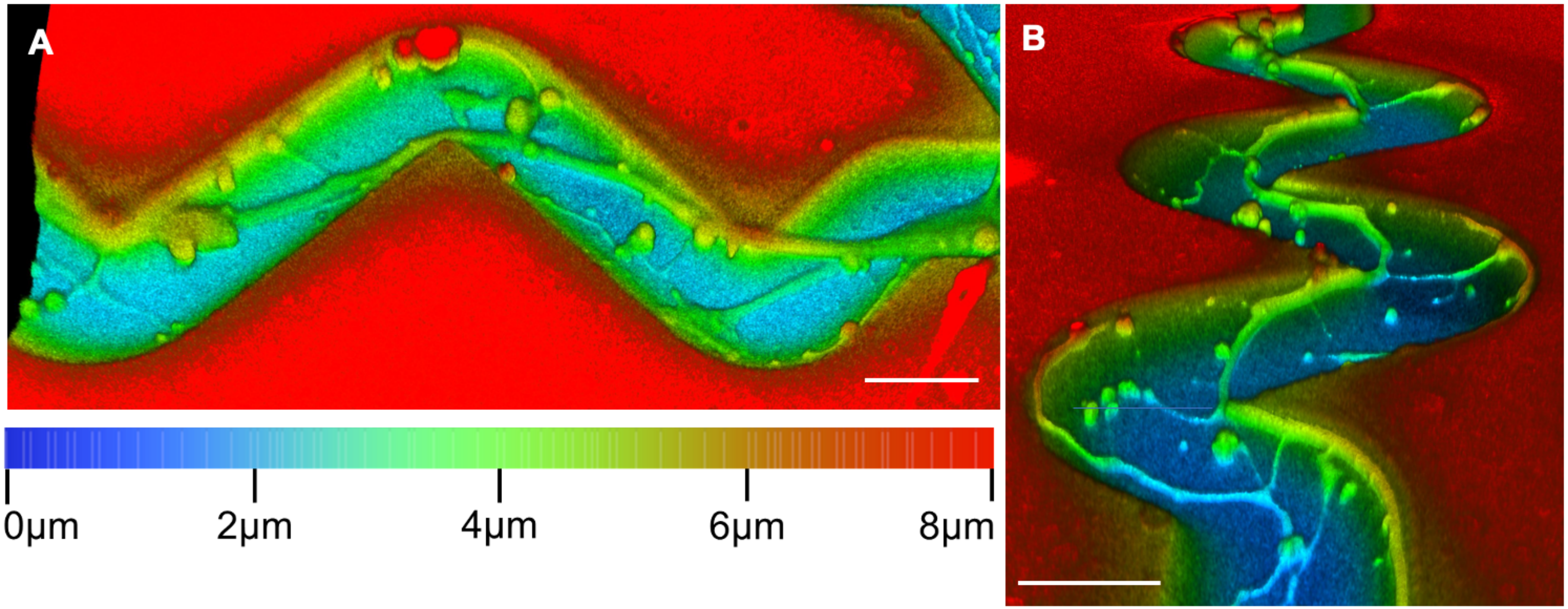

### Supplemental Figure 5

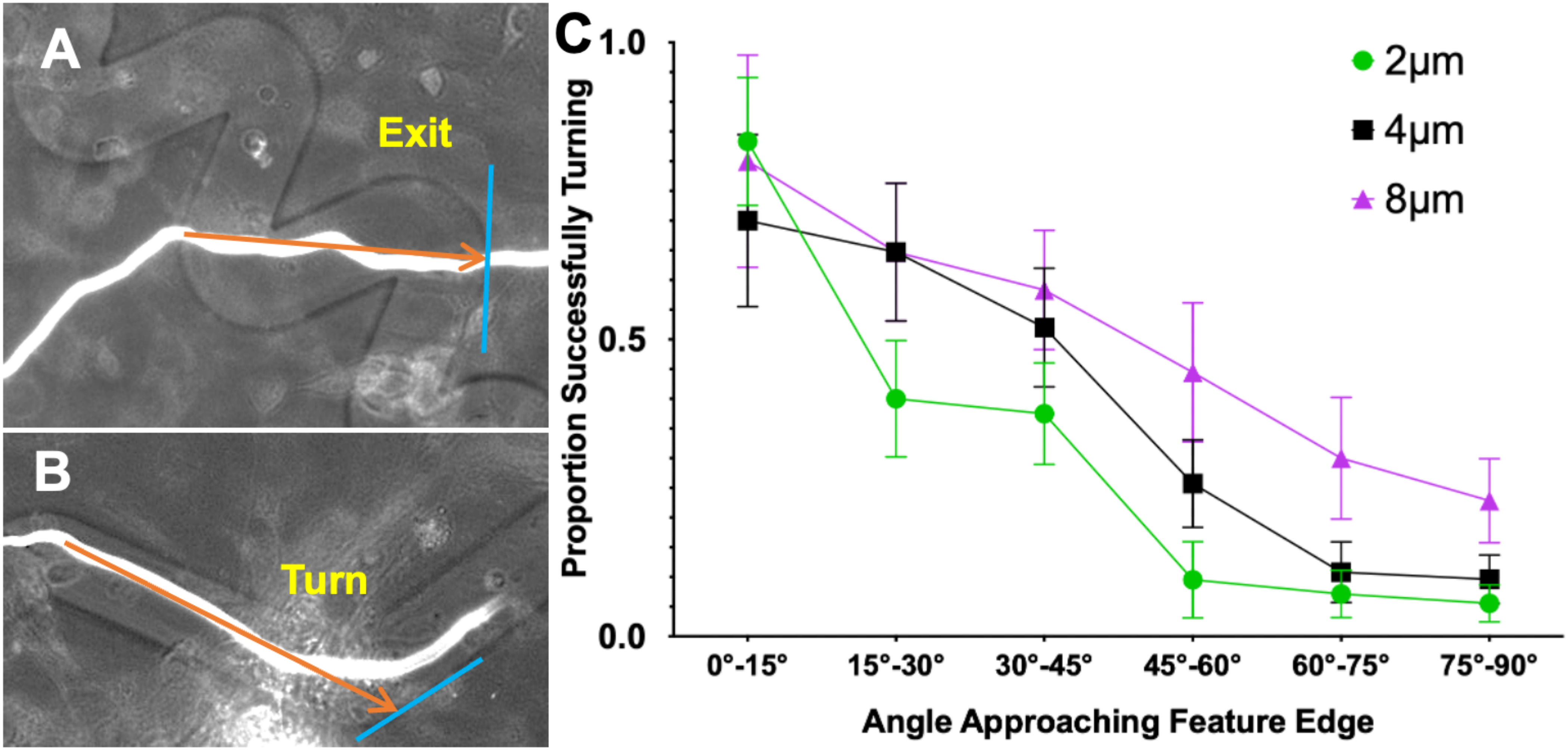
